# A dual role in virion attachment and entry makes the human cytomegalovirus gHgLgO trimer the central player in virion infectivity

**DOI:** 10.1101/2024.11.29.625639

**Authors:** Lena Thiessen, Roberto Garuti, Lucie Kubic, Miwako Kösters, Divya Amarambedu Selvakumar, Thomas Krey, Irene Görzer, Thomas Fröhlich, Barbara Adler

**Affiliations:** Max von Pettenkofer-Institute and Gene Center, Department of Virology, Faculty of Medicine, Ludwig-Maximilians-University (LMU) Munich, Munich, Germany; German Center for Infection Research (DZIF), partner site Munich, Germany; Laboratory for Functional Genome Analysis LAFUGA, Gene Center, Ludwig-Maximilians-University, Munich, Germany; Center of Structural and Cell Biology in Medicine, Institute of Biochemistry, University of Lübeck, Lübeck, Germany; German Center for Infection Research (DZIF), partner site Hamburg-Lübeck-Borstel-Riems, Lübeck, Germany; Institute of Virology, Hannover Medical School, Hannover, Germany; Excellence Cluster 2155 RESIST, Hannover Medical School, Hannover, Germany; Centre for Structural Systems Biology (CSSB), Hamburg, Germany; Center for Virology, Medical University of Vienna, Vienna, Austria

## Abstract

Glycoproteins in the viral envelope of human cytomegalovirus (HCMV) orchestrate virion tethering, receptor recognition and fusion with cellular membranes. The glycoprotein gB acts as fusion protein. The gHgL complexes gHgLgO and gHgLpUL(128,130,131A) define the HCMV cell tropism. Studies with HCMV lacking gO had indicated that gO, independently of binding to its cellular receptor PDGFRα, plays an important second role in infection. Here, we identified a gO mutation which abolished virus particle infectivity by preventing the interaction of gHgLgO with host cell heparan sulfate proteoglycans (HSPGs). We could not only show that gHgLgO – HSPG interactions are a genuine second role of gHgLgO, but also that gHgLgO is the main player in determining the infectivity of HCMV virus particles. This challenges long-accepted textbook knowledge on the role of gB and gMgN complexes in virion tethering. Additionally, it adds the gHgLgO complex to the antigens of interest for future HCMV vaccines or treatments.

## Introduction

Infections with human cytomegalovirus (HCMV) are widespread in the human population and persist like all herpesviruses for life^1^. In immunocompromised individuals, infections can result in life-threatening diseases and infections of the fetus are a major cause of congenital birth defects^1^. Therefore, significant efforts have been made to develop vaccines for preventing or restricting infection mainly by using virion glycoprotein complexes as antigens^2–4^. Up to now, none of the vaccines has been highly efficient.

Infection of target cells by HCMV involves three steps: tethering to cell surfaces, receptor recognition and fusion with cellular membranes^5,6^. Several virion glycoprotein complexes are involved in these steps including the trimeric gHgLgO complex, the pentameric gHgLpUL(128,130,131A) complex and the gB homotrimer. The heterodimer gHgL and the gB trimer form the viral fusion machinery while the gHgL-associated proteins promote binding to host cell receptors^5,6^. Deletion of the gHgL-associated proteins of the pentamer results in restriction of the HCMV cell tropism^5^. In contrast, deletion of gO results in a more dramatic reduction of infectivity for all host cells^7–13^. The cellular entry receptor bound by gHgLgO is platelet-derived growth factor receptor alpha (PDGFRα)^11,12,14–16^, yet loss of this interaction cannot explain the loss of infectivity for PDGFRα-negative cells. Several studies addressing impaired infectivity of HCMV mutants lacking gO had pointed towards a second role of gHgLgO in initiating cell infection^13,14,17,18^. When studying the role of the functionally homologous gHgLgO complex of murine cytomegalovirus (MCMV) *in vivo*, we have shown that deletion of gO nearly completely abolished susceptibility of mice to MCMV infection by preventing infection of different first target cells^19^. In contrast, deletion of the MCMV gHgLMCK2 complex, which is functionally homologous to the pentameric complex of HCMV, only resulted in a minor impairment of infection^19–22^. Thus, the gHgLgO complex might be an interesting vaccine target. Yet, for the development of vaccines targeting the HCMV trimer, a prerequisite would be to fully understand the role of gHgLgO in infection.

Here, we used an HCMV gO mutant in which five basic amino acids in the highly conserved peptide site 249 to 254 of gO were exchanged to alanines^13^. This mutant showed a strong and cell-type independent loss of infectivity but normal binding to PDGFRα. A profound analysis of this mutant revealed the postulated, but until now unidentified second role of gHgLgO in HCMV infection. We could show that tethering of HCMV particles to host cells is nearly exclusively driven by the interaction of the gHgLgO complex with heparan sulfate proteoglycans (HSPGs), a function whose loss goes along with the loss of particle infectivity. We could also show that complexes of gHgLgO and gB^14^ do not contribute to HSPG binding. Complex formation of gHgLgO with gB could even block the intrinsic capacity of gB^23^ to bind HSPGs.

## Results

### HCMV mutants revealing the dual role of glycoprotein gO

Deletion of the HCMV glycoprotein gO results in elimination of PDGFRα-dependent HCMV entry^11,12,14,24^. In addition, gO-negative HCMV particles show a dramatic reduction in infectivity for all host cells which is independent of PDGFRα^10,13,14,17^. So far, all attempts to understand this second role of gO were unsuccessful. We screened the literature for HCMV gO mutants, which exhibited a major loss of particle infectivity for different host cells, but were still capable of forming gHgLgO complexes in virions. A mutagenesis of conserved regions in gO described a mutant with the desired phenotype^13^. This mutant comprised a change of amino acids 249-**RK**L**KRK**-254 to 249-**AA**L**AAA**-254 and will here be called gO249. For a thorough characterization of gO249, we compared it with a second gO mutant with amino acids 117-**RK**PA**K**-121 changed to 117-**AA**PA**A**-121^25^, here called gO117. The gO117 mutant has been shown to be impaired in binding to PDGFRα. Based on recently published cryo EM structures of the trimer^17^, the gO249 mutation could be localized proximal to glycoprotein gL and opposite to the PDGFRα binding site (Fig. 1a). The gO117 mutation could be localized to the PDGFRα interface (Fig. 1a). We introduced these mutations into TB40-BAC4-luc virus^8^ (here designated as WT virus). Western blot (WB) analysis of virus particle lysates revealed comparable contents of the virion glycoproteins gH, gO and gB for WT, gO249 and gO117 (Fig. 1b, upper panels). Under non-reducing conditions, the quantities and electromobilities of gHgLgO complexes of gO249 and gO117 were comparable to those of WT virus (Fig. 1b, lower panels). For comparison, extracts of gO-negative virions (ΔgO mutant)^8^ are shown (Fig. 1b). To compare the infection capacities of WT, gO249, gO117 and ΔgO viruses for fibroblasts and endothelial cells, PDGFRα-positive human foreskin fibroblasts (HFF) and PDGFRα-negative immortalized microvascular endothelial cells (TIME cells) were chosen (Supplementary Fig. 1a). HCMV entry into TIME cells is strictly dependent on the pentameric complex^8,14,26^, whereas entry into HFF predominantly depends on the trimeric gHgLgO complex^14^. Cells were infected with equal numbers of virus particles and infection capacities evaluated using a luciferase assay (Fig. 1c). Consistent with its first characterization^13^, gO249 showed strongly reduced infection capacities for fibroblasts (0.3% of WT) and endothelial cells (0.2% of WT). Compared with gO249, infection capacities of gO117 were less reduced (HFF: 13.4% of WT; TIME cells: 12% of WT). Infection capacities of the ΔgO mutant showed a reduction to 0.01% on HFF and 1.7% on TIME cells when compared to WT virus. The stronger reduction of ΔgO compared to gO249 on HFF may reflect the loss of additional gO functions, whereas the reduced attenuation on TIME cells might reflect elevated levels of pentameric complex in gO-negative virions as described before^14,27–29^.

**Figure 1.**
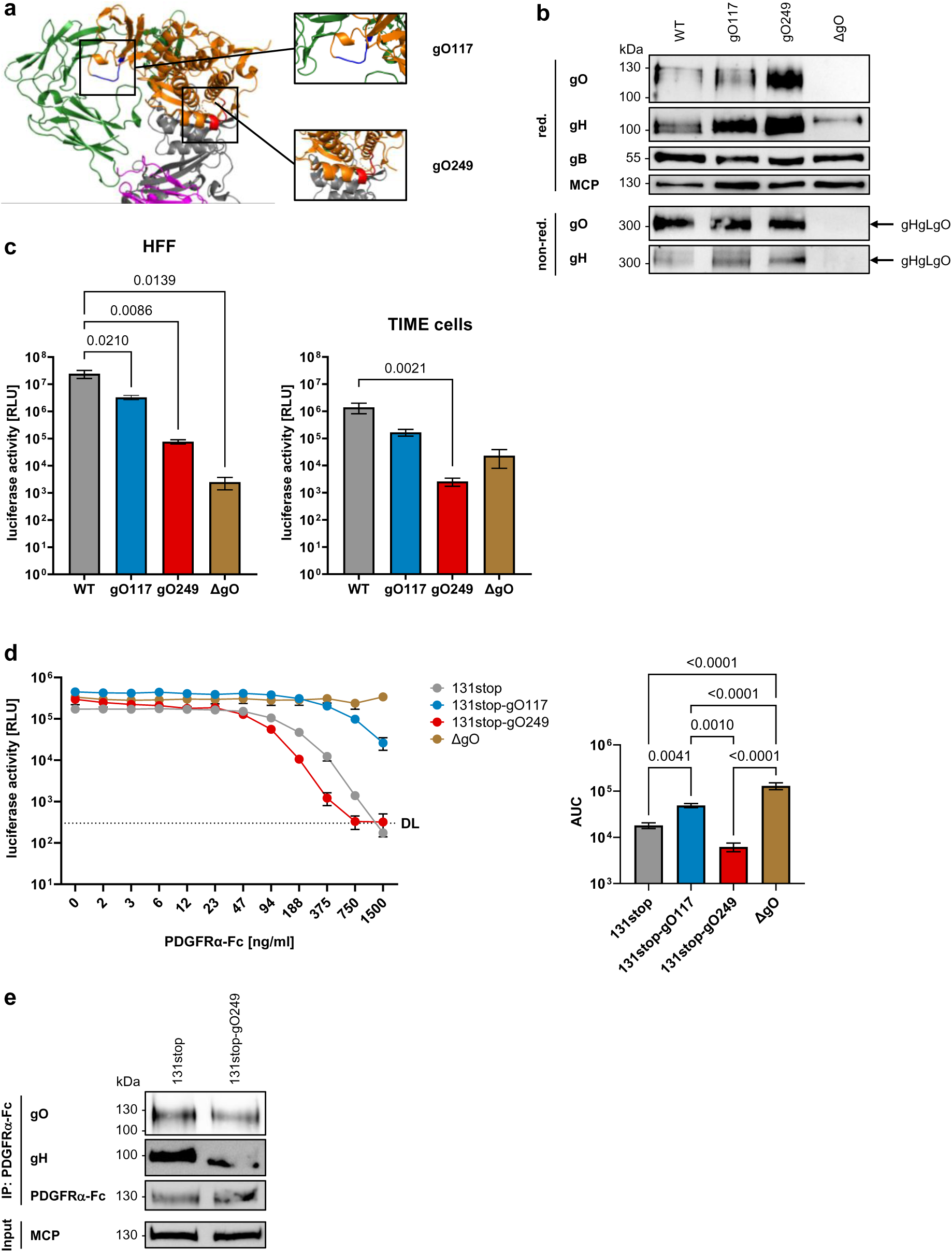
The gO249 mutation results in a PDGFRα-independent loss of particle infectivity. (**a**) Cryo EM structure^16^ of gHgLgO bound to PDGFRα (gH (purple), gL (grey), gO (orange), PDGFRα (green); PDB: 7LBF) with locations of mutated sequences of gO117 and gO249 highlighted in blue and red, respectively. (**b**) Levels of virion glycoproteins gH, gO and gB were assessed in virion lysates by WB analysis under reducing (red.) conditions. The formation of gHgLgO complexes was assessed under non-reducing (non-red.) conditions. Levels of major capsid protein (MCP) reflect comparable numbers of virus particles. One representative experiment of three is shown. (**c**) HFF (left panel) or TIME cells (right panel) were infected with 100 virus particles/cell and infection was assessed by luciferase assay 48 h p.i. Shown are means +/− SEM of independent experiments (*n* = 3-6). *P* values of statistically significant differences are depicted (HFF: one-way ANOVA; TIME cells: Kruskal-Wallis test). (**d**) Infection of HFF after preincubation of viruses with increasing concentrations of soluble recombinant PDGFRα-Fc. 48 h p.i., infection was assessed by luciferase assay. Shown are means +/− SEM of independent experiments (*n* = 3-6). Right panel: Calculation of the area under the curve (AUC) of normalized data (one-way ANOVA). *P* values of statistically significant differences are depicted. (**e**) WB analysis of gH and gO precipitated from virion lysates using soluble recombinant PDGFRα-Fc. One representative experiment of three is shown. RLU: relative light units. DL: detection limit.

The differences in infection capacities also shaped the growth curves of gO mutants (Supplementary Fig. 1b). HFF were infected with equal infectious doses of WT, gO249, gO117 or ΔgO virus and virus growth was monitored by determining infectious virus in cell culture supernatants. Particle numbers and specific infectivity, exemplarily determined for day eight supernatants, confirmed that the gO249 mutation or deletion of gO impaired particle infectivity, but not particle release. The reduced particle infectivity became apparent in the growth curves by a strongly reduced production of infectious virus.

To evaluate binding of the gHgLgO complex of the gO249 mutant to the cellular PDGFRα receptor, we infected HFF with WT, gO249, gO117 and ΔgO viruses pre-incubated with increasing concentrations of soluble recombinant PDGFRα-Fc protein. With the exception of ΔgO, a pentamer-negative genomic background (TB40-BAC4-luc-131stop virus^8^, here called 131stop virus) was chosen to restrict infection of WT, gO249 and gO117 viruses to gHgLgO-driven entry. As shown before^14^, 131stop virus could be completely inhibited with high concentrations of recombinant PDGFRα protein, whereas the pentamer-dependent infection with ΔgO virus could not be inhibited at all (Fig. 1d). The 131stop viruses expressing gO_WT_ or gO_249_ were inhibited to a similar extent which suggested that the gO249 mutation does not interfere with PDGFRα binding. In contrast, gO117 virus was inhibited to a much lesser extent. Equal binding of gHgLgO_WT_ and gHgLgO_249_ to PDGFRα could be confirmed by immunoprecipitation experiments using recombinant PDGFRα-Fc to precipitate the complexes from virion lysates (Fig. 1e) or from lysates of HEK293T cells transfected with plasmids expressing gH, gL, gO_WT_ or gO_249_ (Supplementary Fig. 1c).

### gHgLgO shapes HCMV particle infectivity by promoting attachment to host cells

Several studies had suggested or shown that gHgLgO is crucial for particle infectivity independent of PDGFRα and the host cell^10,13,17,18^. The gO249 mutant has lost most of its particle infectivity without a change in PDGFRα-binding. This phenotype implied a widespread second cell-surface (co-)receptor for gHgLgO. Therefore, we searched for surface proteins of HFF which can bind virions containing gO_WT_, but not virions containing gO_249_. We used 131stop and 131stop-gO249 viruses to exclude interactions of surface proteins with the pentameric complex. We co-incubated HFF with equal numbers of virus particles, lysed cells and viruses bound to cell surfaces and precipitated gHgLgO complexes from these lysates using a gH-specific antibody followed by mass spectrometry (LC-MS/MS) of the precipitated proteins (Fig. 2a). The anti-gH antibody used equally precipitated gHgLgO_WT_ and gHgLgO_249_ complexes (Fig. 2b). When comparing proteins co-precipitated from HFF-bound 131stop or 131stop-gO249 virions, 51 hits were identified which were significantly lower for the 131stop-gO249 mutant (Fig. 2c and Supplementary Table 1). These hits included the glycoproteins of the gHgLgO complex, the gHgLgO-associated glycoprotein gB^14^, known HCMV particle-associated proteins like exosome-associated proteins^30^, the gHgLgO cellular receptors PDGFRα and TGFβRIII^16^ and cell surface heparan sulfate proteoglycans (HSPGs) like syndecans and glypican. The low levels of gH, gL, gO and gB found for 131stop-gO249 suggested that binding of 131stop-gO249 virions to HFF was impaired. To address this, we repeated the experiments, but this time quantified virions bound to cell surfaces using qPCR. Binding of 131stop virions containing gO_249_ was strongly reduced when compared to virions containing gO_WT_ (Fig. 2d) which confirmed that gO249 virions exhibit a defect in cell attachment. We observed this difference also with WT and gO249 viruses expressing additionally the pentameric complex (Fig. 2e) and when endothelial cells were used as host cell targets (Fig. 2f). In summary, these data suggest that gO strongly contributes to virion tethering through a PDGFRα-independent interaction with host cell surfaces.

**Figure 2.**
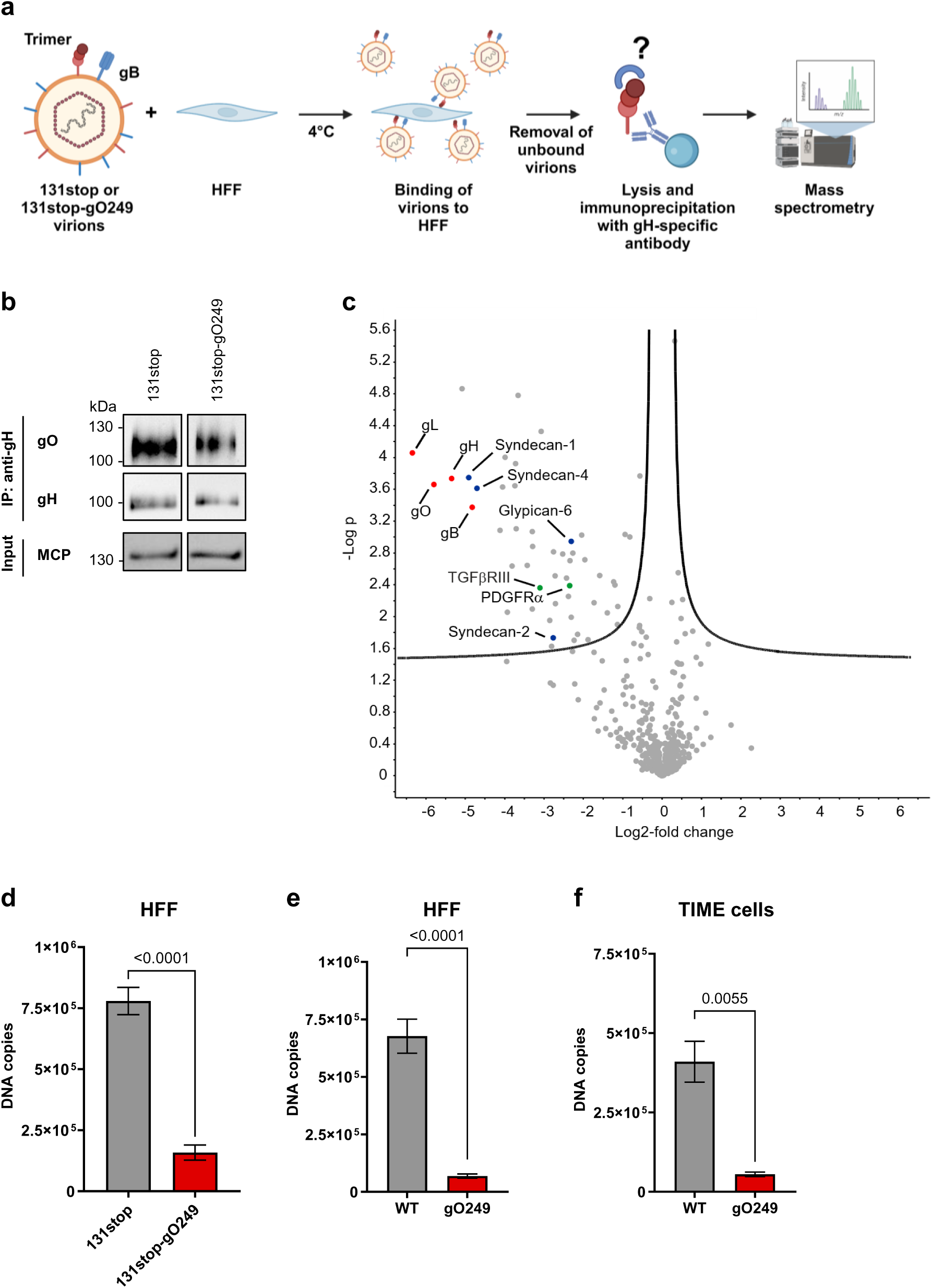
Screening for gHgLgO interaction partners reveals decreased binding of the gO249 mutant to cell surfaces. (**a**) Schematic presentation of experiments performed to identify cell surface proteins which interact with gHgLgO_WT_ or gHgLgO_249_. 131stop or 131stop-gO249 virions were co-incubated with HFF for 1 h at 4°C. Unbound virions were removed and cells were lysed subsequently. An immunoprecipitation was performed using the gH-specific antibody 14-4b followed by LC-MS/MS analysis to identify protein interaction partners of gH for 131stop (gO_WT_) and 131stop-gO249 (gO_249_) viruses. (**b**) WB analysis of gH and gO immunoprecipitated from lysates of 131stop and 131stop-gO249 virions using the gH-specific antibody 14-4b. One representative experiment of four is shown. (**c**) Volcano plot of LC-MS/MS data from anti-gH immunoprecipitates of lysates of HFF co-incubated with 131stop or 131stop-gO249 virions as described under (a). Data from 131stop-gO249 virions were compared to 131stop virions and depicted as –Log *p*-value versus Log2-fold change. HCMV glycoproteins are highlighted in red, the gHgLgO receptors PDGFRα and TGFβRIII in green and HSPGs in blue. Data are derived from three independent experiments. (**d**) HFF were incubated with 5×10^7^ 131stop or 131stop-gO249 virus particles and bound particles were quantified by qPCR. Shown are means +/− SEM of five independent experiments. (**e**) HFF or (**f**) TIME cells were incubated with 5×10^7^ particles of WT or gO249 virus particles and bound particles were quantified by qPCR. Shown are means +/− SEM of independent experiments (*n* = 3-6). (**d-f**) *P* values of statistically significant differences are depicted (Student’s unpaired t-test).

### gHgLgO - HSPG interactions are major determinants of HCMV particle infectivity

The strongly reduced infectivity and binding capacity of gO249 virus particles suggested that the gO249 mutation abolished the interaction of virions with cell surface molecules promoting virus attachment. Interestingly, four of the proteins co-precipitated with gHgLgO were heparan sulfate proteoglycans (HSPGs). Adsorption of virus particles to cell surface HSPGs is an accepted first step in herpesvirus and HCMV infection^31,32^. The gO249 mutation (249-**RK**L**KRK**-254 to 249-**AA**L**AAA**-254) reverted a linear sequence of basic amino acids, a motif characteristic for HSPG-binding proteins^33^, into a non-polar amino acid sequence. Therefore, we hypothesized that gHgLgO may directly or indirectly interact with HSPGs and that the reduced cell surface binding of gO249 may be due to an impaired interaction of gHgLgO_249_ with HSPGs. To address this, we infected HFF and TIME cells with equal infectious doses of viruses expressing gO_WT_ or gO_249_ and for comparison also with ΔgO virus in the presence of increasing concentrations of the HSPG mimic heparin. While gO_WT_ viruses could be inhibited in a concentration-dependent manner, both on HFF and TIME cells, infections with gO_249_ and ΔgO viruses were resistant to heparin (Fig. 3a and b). This indicated that gO and more specifically amino acids 249 to 254 of gO promote the virion - HSPG interaction. HSPG-dependence of gO_WT_ virus infection and no HSPG-dependence of gO_249_ viruses could also be observed when host cells were pre-treated with heparinase (Fig. 3c and d).

**Figure 3.**
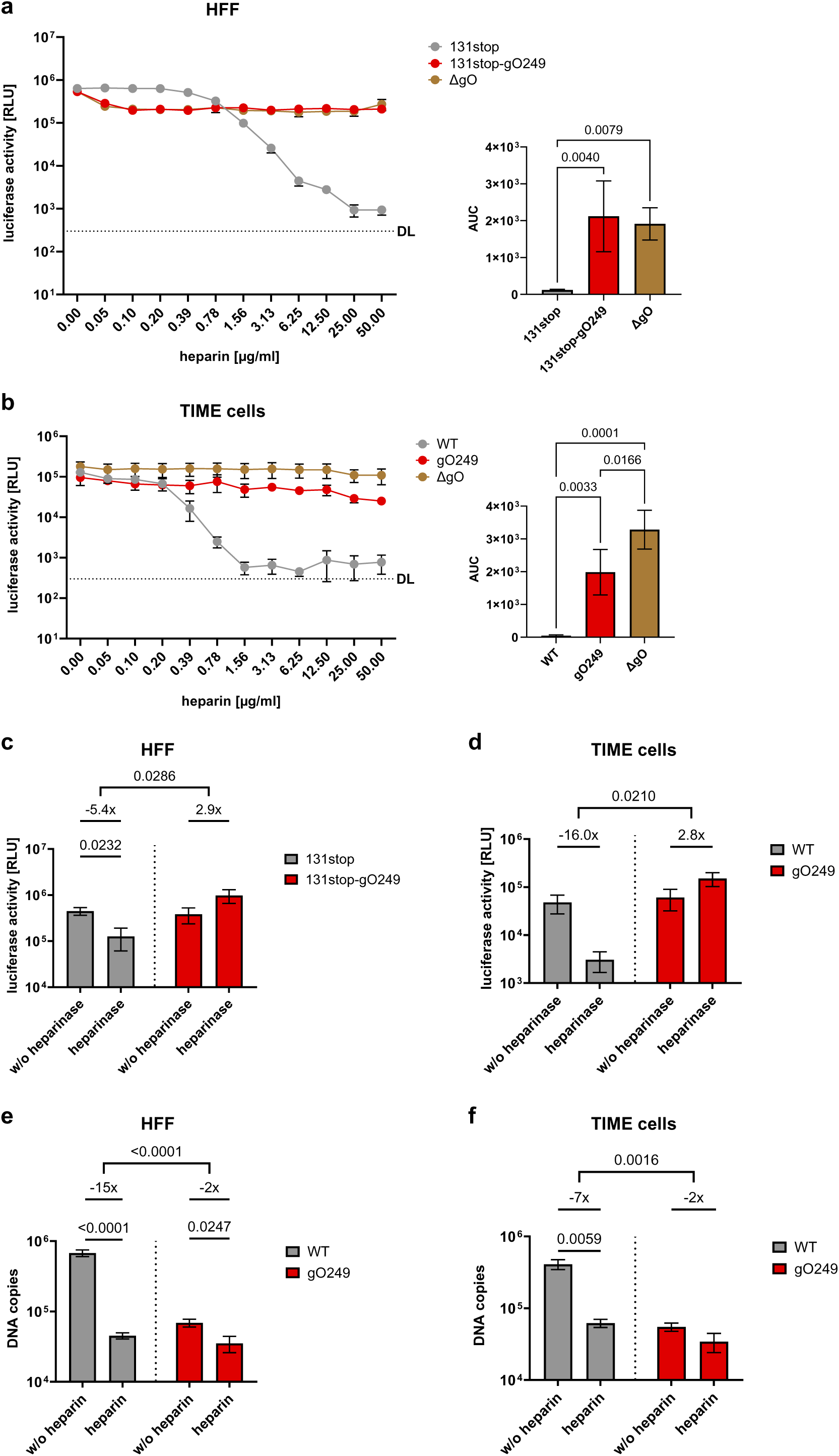
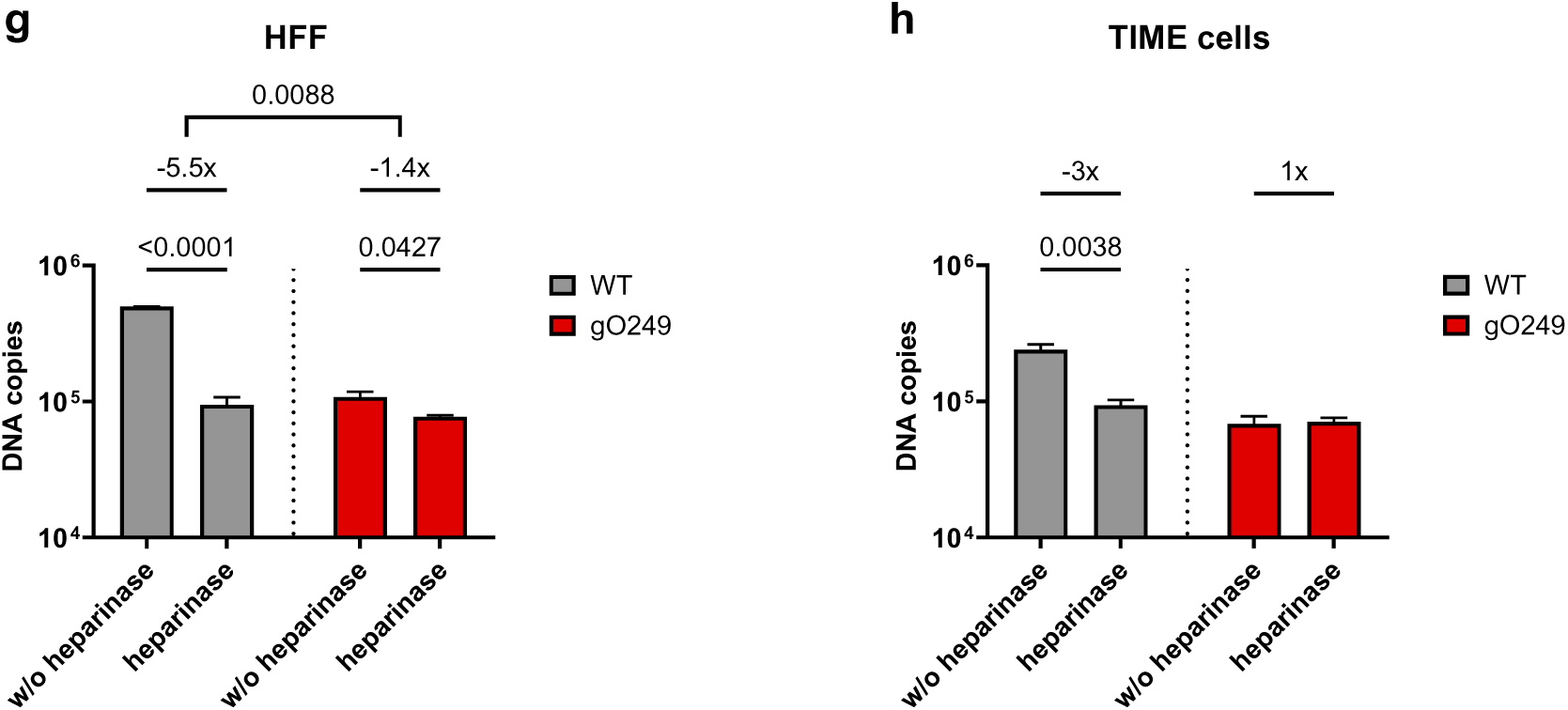
Infection and virion attachment require interaction of gHgLgO with HSPGs. (**a**) HFF and (**b**) TIME cells were infected with 131stop, 131stop-gO249 or ΔgO virus in the presence of increasing concentrations of heparin. 48 h p.i., infection was assessed by luciferase assay. Shown are means +/− SEM of three independent experiments. Right panels: Calculation of the AUC of normalized data (one-way ANOVA). (**c**) HFF or (**d**) TIME cells were pre-treated with heparinase I, II and III followed by infection with 131stop or 131stop-gO249 virus (HFF) or WT or gO249 virus (TIME cells). 48 h p.i., infection was assessed by luciferase activity. Shown are means +/− SEM of independent experiments (*n* = 3-4). Statistical significance was determined for pairwise comparisons of heparinase-treated or -untreated cells (Student’s unpaired t-test or Mann-Whitney test). Additionally, fold changes of pairwise comparisons were calculated and analyzed (Student’s unpaired t-test or Mann-Whitney test). (**e** and **g**) HFF or (**f** and **h**) TIME cells were incubated with 5×10^7^ WT or gO249 virus particles in the presence (100 µg/ml) or absence of heparin (**e** and **f**) or after pre-treatment of the cells with heparinase I, II and III (**g** and **h**) and virus particles bound to cells were quantified by qPCR. Shown are means +/− SEM of independent experiments (*n* = 3-6). Statistical significance was determined for pairwise comparisons of treated or -untreated infections (Student’s unpaired t-test). Additionally, fold changes of pairwise comparisons were calculated and analyzed (Student’s unpaired t-test and Mann-Whitney test). The data showing virus binding in the absence of heparin (**e** and **f**) are identical to the data of Figure 2e and f. (**a-h**) *P* values of statistically significant differences are depicted. DL: detection limit.

Differences in HSPG interactions between WT and gO249 viruses should also become apparent when binding to cells is studied. Binding of WT virus to HFF and TIME cells was clearly inhibited in the presence of heparin (Fig. 3e and f) and by heparinase pre-treatment of the host cells (Fig. 3g and h). Binding of gO249 virus could not much further be inhibited (Fig. 3e to h). Thus, the 249 mutation of gO strongly impaired HSPG-dependent tethering of virions which provides a highly plausible explanation for the low infectivity of gO249 virus particles.

We also compared binding of gHgLgO_WT_ and gHgLgO_249_ complexes to heparin by precipitating gHgLgO from 131stop and 131stop-gO249 virion lysates using heparin agarose (Fig. 4a). Precipitation of gHgLgO_249_ was strongly reduced and went along with reduced precipitation of gB from 131stop-gO249 virion lysates. Thus, very likely, virion gB alone does not efficiently bind to heparin, but is co-precipitated as it forms a complex with gHgLgO^14^. This raised the question whether gHgLgO - gB complexes are required for HSPG binding and whether the impaired binding of gHgLgO_249_ to heparin binding reflects the loss of gHgLgO - gB complex formation. To address the latter point, we compared the interaction of gB with gHgLgO_WT_ and gHgLgO_249_ by co-precipitating gH, gO and gB from virion lysates using recombinant PDGFRα-Fc or a gH-specific antibody. The interaction of gHgLgO_249_ with gB was not reduced, but even 4-fold stronger than the interaction of gHgLgO_WT_ and gB (Fig. 4b). Yet, we could not exclude that the gHgLgO_249_ – gB complex is specifically impaired in binding to HSPG. To exclude that gHgLgO needs gB to bind to HSPG, we expressed recombinant gH, gL and gO in HEK293T cells. Both, gO_WT_ and gO_249_ formed gHgLgO complexes in transfected cells (Fig. 4c). WB analyses of lysates of infected HFF showed two distinct bands for gO_WT_ and gO_249_ of whom only the upper bands could be detected in lysed virions (Supplementary Fig. 2). These double bands were also detectable in transfected cells (Fig. 4d, left panel). When using heparin agarose to precipitate the recombinant gHgLgO complexes, we found that the upper gO band could only be precipitated from cells expressing recombinant gHgLgO_WT_ and not from cells expressing gHgLgO_249_ (Fig. 4d, right panel). This indicated that the gHgLgO complex alone is sufficient to bind to heparin and that the gO249 mutation abolishes this interaction.

**Figure 4.**
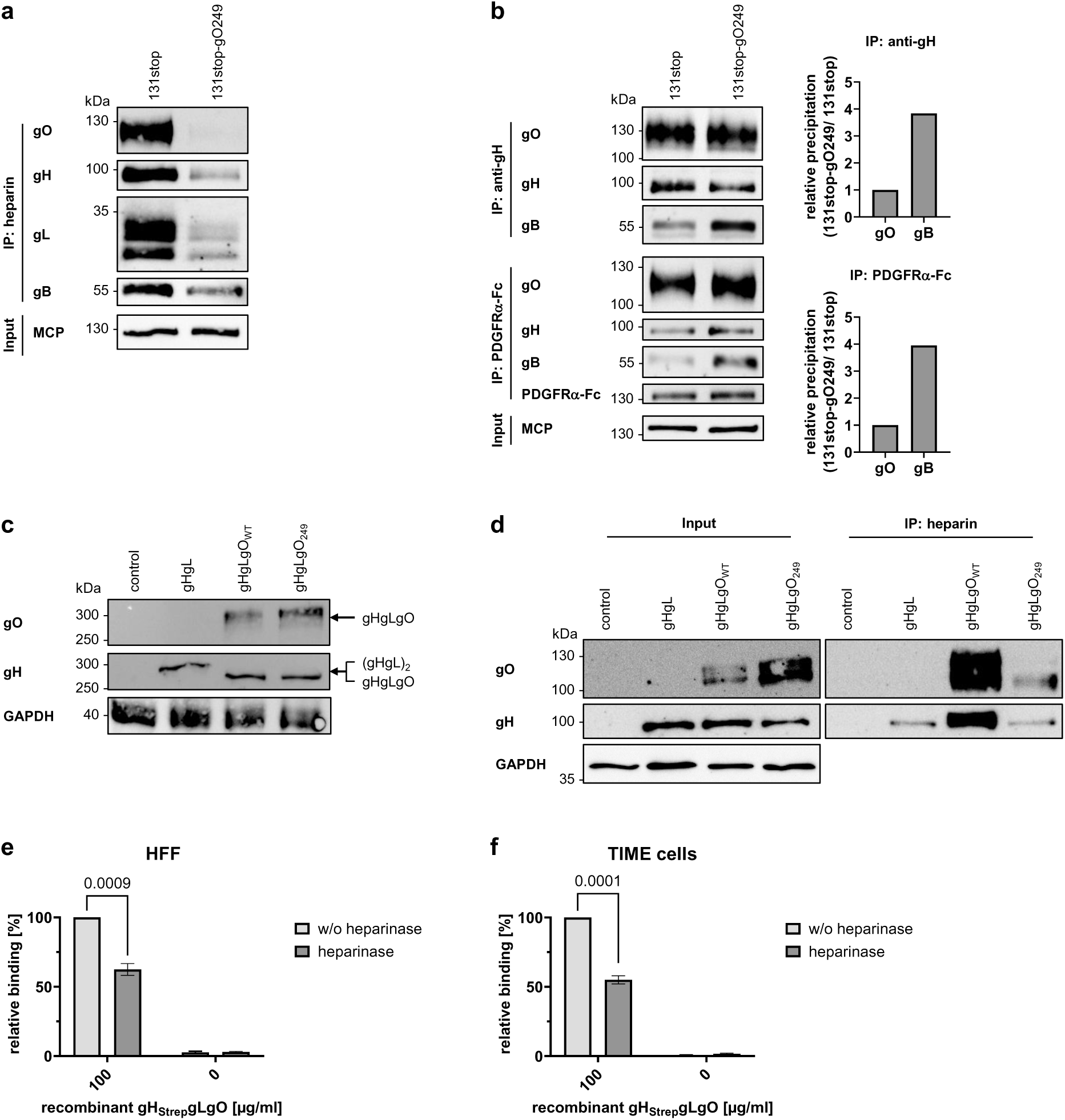
gHgLgO is sufficient to bind to HSPGs and mutation of amino acids 249 to 254 of gO abolishes this interaction. (**a**) WB analysis of gH, gL, gO and gB precipitated from 131stop and 131stop-gO249 virion lysates using heparin agarose. One representative experiment of four is shown. (**b**) WB analysis of gH, gO and gB precipitated from 131stop and 131stop-gO249 virion lysates using an anti-gH antibody (14-4b) or soluble recombinant PDGFRα-Fc. Representative experiments of three are shown. Right panels: ImageJ quantification of gO and gB WB signals. gO precipitation levels were normalized and relative gB levels determined. (**c**) WB analysis of gH and gO expression in lysates of HEK293T cells transfected with plasmids expressing GFP (control), gH and gL (gHgL), gH, gL and gO_WT_ (gHgLgO_WT_) or gH, gL and gO_249_ (gHgLgO_249_). Lysates were prepared under non-reducing conditions. GAPDH levels served as loading control. One representative experiment is shown. (**d**) WB analysis of gH and gO precipitated from lysates of HEK293T cells expressing GFP (control), gHgL, gHgLgO_WT_ or gHgLgO_249_ using heparin agarose. GAPDH levels served as loading control. One representative experiment is shown. (**e**) HFF and (**f**) TIME cells were pre-treated with heparinase I, II and III followed by incubation with recombinant gH_Strep_gLgO_WT_ (100 µg/ml). Binding of recombinant protein to cells was assessed using an ELISA detecting Strep-tagged proteins. Binding of recombinant gH_Strep_gLgO_WT_ to untreated cells was arbitrarily set to 100%. Shown are means +/− SEM of three independent experiments. Statistical significance was determined by pairwise comparison of heparinase-treated and -untreated cells (Student’s unpaired t-test). *P* values of statistically significant differences are depicted.

To confirm the interaction of gHgLgO with HSPG in a more physiological setting, namely on the surface of the host cells, we co-incubated heparinase-treated and untreated HFF and TIME cells with soluble recombinant gH_Strep_gLgO_WT_ complex and measured binding of this complex in an ELISA detecting the Strep-Tag of gH. We observed a significant reduction of binding of gHgLgO to heparinase treated HFF and TIME cells (Fig. 4e and f). Thus, attachment of gHgLgO to cell surfaces is dependent on an interaction with HSPG and is not restricted to cells expressing PDGFRα.

We could hardly precipitate gHgLgO_249_ complexes using heparin, but still observed residual gB precipitation from gO249 virion lysates and residual inhibition of gO249 binding. Therefore, we wondered whether ΔgO virions lacking gHgLgO complexes show a complete loss gB precipitation by heparin agarose. Interestingly, gB could be precipitated from ΔgO virions indicating a gB - heparin interaction independent of gHgLgO (Supplementary Fig. 3a). When comparing virion binding of WT, gO249 and ΔgO viruses to HFF and TIME cells, we observed an enhanced binding of ΔgO virions compared to gO249 virions which could be completely inhibited by heparin (Supplementary Fig. 3b and c). In summary, these data might be interpreted such that gHgLgO is the major driver of virion attachment by binding to HSPGs. gB can also bind HSPGs, although to a much lesser extent. The capacity of gB to bind HSPGs becomes particularly apparent in ΔgO infections. The gO249 mutation not only results in a loss of the gHgLgO - HSPG interaction, but very likely additionally blocks binding of gB to HSPGs by trapping gB in tight gHgLgO_249_ - gB complexes.

### gHgLgO - from tethering to cell surface HSPGs to PDGFRα docking

Infectivity of cell free HCMV is shaped by binding of the gHgLgO trimer to host cell surface HSPGs and docking of the trimer or pentamer to HCMV entry receptors like PDGFRα or NRP2^34^, respectively. Assuming that gHgLgO binding to HSPGs is the first step in cell surface binding, this interaction with a relatively abundant cell surface molecule^31^ either has to leave room for an additional interaction with the entry receptor PDGFRα or, if binding is overlapping, PDGFRα-interactions have to successfully compete with HSPG interactions. We tested this by consecutive addition of heparin and PDGFRα in immunoprecipitation experiments using virion lysates. Addition of recombinant PDGFRα-Fc to heparin agarose beads released most of gHgLgO and gB bound to the beads (Fig. 5a). In contrast, gHgLgO and gB bound to PDGFRα-Fc on sepharose beads could not be removed by addition of heparin (Fig. 5b). This indicated that binding to HSPGs and PDGFRα is competitive and that binding of gHgLgO to PDGFRα is stronger than binding of gHgLgO to heparin. We suggest a model for cell free infection in which gHgLgO determines efficient tethering of virus particles to cell surfaces followed by firm docking to entry receptors (Fig. 6).

**Figure 5.**
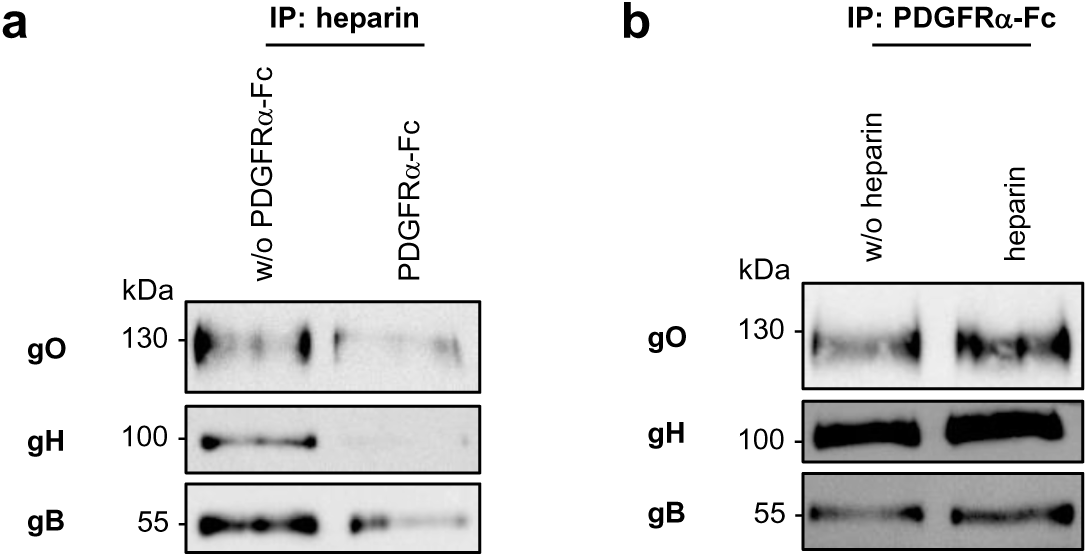
Competitive binding of heparin and PDGFRα to gHgLgO suggests overlapping binding sites. gHgLgO complexes were precipitated from lysates of 131stop virions using (**a**) heparin agarose or (**b**) soluble recombinant PDGFRα-Fc. (**a**) Proteins bound to heparin agarose beads were then co-incubated with PDGFRα-Fc (4 µg/ml) for 4 h at 4°C. (**b**) Proteins bound to PDGFRα-Fc/Protein A sepharose beads were co-incubated with heparin (500 µg/ml) for 4 h at 4°C. (**a, b**) Afterwards, beads were washed, incubated with 2x sample buffer and analyzed by WB for gH, gO and gB. Shown are representative experiments.

**Figure 6.**
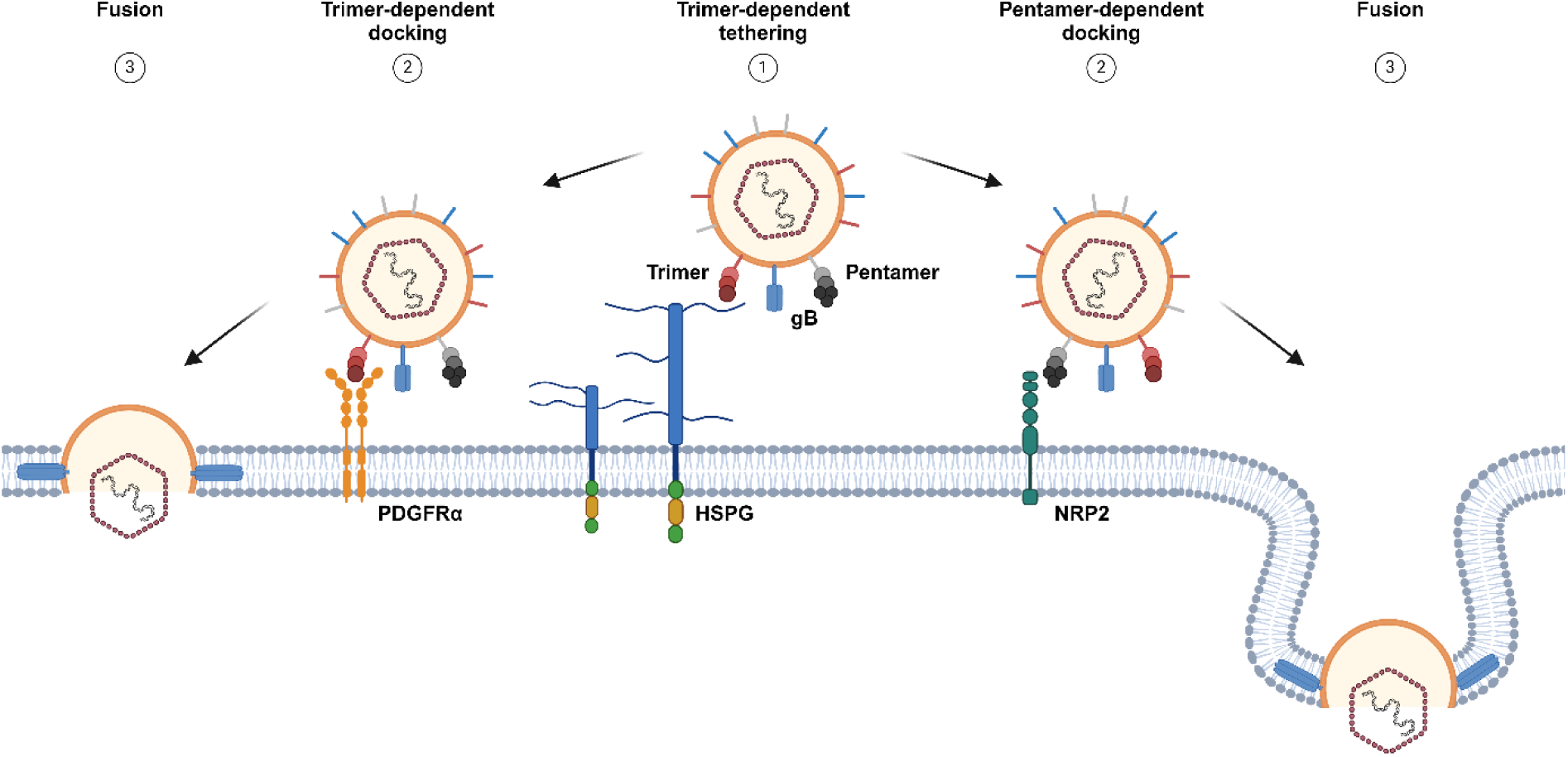
A model for the role of the HCMV trimer in infection of cells with free virus. In a first step, trimer – HSPG interactions tether virus particles to cell surfaces (1). gHgLgO-specific tethering is followed by docking of trimer or pentamer to their specific host cell entry receptors (2) and fusion of the viral envelope with cellular membranes (3). Exemplarily, the trimer-specific receptor PDGFRα and the pentamer-specific receptor NRP2 are shown.

## Discussion

Cytomegaloviruses express two alternative gHgL glycoprotein complexes, gHgLgO and gHgLpUL(128,130,131A) which promote receptor recognition. For both, several host cell proteins have been described as binding partners and entry receptors^16,34,35^. HCMV can still infect cells when only one of the alternative gHgL complexes is deleted, yet the outcomes of deletion of the trimer or the pentamer are completely different. In cell culture, deletion of the pentamer abolishes infection of cell types like endothelial and epithelial cells, macrophages and dendritic cells, but for example not of fibroblasts^36^. In contrast, deletion of the trimer results in a drastic loss of infectivity for all host cells^9,10^. The gHgLgO complex uses PDGFRα as an entry receptor, an interaction which is resolved on a structural basis and also on the level of its role in infection^11,14,16^. Yet, the loss of the gHgLgO - PDGFRα interaction could not explain the cell type-independent loss of infectivity when gO is deleted. gHgLgO can also interact with TGFβRIII^16^ but this cell surface receptor has been excluded as an entry receptor^34^.

Two recent studies had shown that soluble gHgLgO and also gO peptides bind to cell surfaces of PDGFRα-positive and -negative cells and interfere with HCMV infection^17,18^. They postulated additional widespread cell surface receptor(s) promoting a gHgLgO-dependent initial attachment to cells followed by entry receptor docking. Interestingly, their experiments, which were based on staining cell surface-bound trimer excluded an involvement of heparin sulfate moieties^17^.

Instead of comparing binding of recombinant gHgLgO complexes or comparing gO-positive or -negative virions, we wanted to identify these postulated receptor(s) by studying gO mutants with a general loss of virus infectivity, but not PDGFRα binding. A common strategy to identify mutants of interest is a protein-wide mutagenesis. Recently, a mutagenesis of conserved sites of gO has been published^13^. From this publication, we chose the gO249 mutant with amino acid sequence 249-**RK**L**KRK** -254 mutated to 249-**AA**L**AAA**-254. The gO249 mutant was still able to form virion gHgLgO complexes, but showed a drastic loss of infectivity for PDGFRα−positive and -negative cells, just like gO-negative HCMV. Here, we could complement the initial characterization^13^ by showing that gO249 virions interact with PDGFRα like WT virions. We chose a quantitative mass spectrometry approach to compare cell surface receptor(s) bound by gHgLgO_WT_ or gHgLgO_249_ complexes. We made two important findings. First, gHgLgO_WT_ complexes strongly interact with HSPGs and second, the 249-**RK**L**KRK** -254 site of gO is central for virus attachment to cells. A profound analysis of the gO249 mutant allowed us to identify the gHgLgO complex as the major player in HSPG-driven tethering of virions to cells. Thus, the low infectivity of the gO249 mutant was due to a loss of virion – HSPG interactions. The mutation comprised a linear sequence of basic amino acids, typical for HSPG-binding proteins^33^ and also described for other HSPG-binding herpesvirus glycoproteins like HSV-1 gB^37^. Our findings matched recent studies showing that preincubation of HCMV host cells with gO peptides comprising the 249 to 254 amino acid sequence and pre-incubation of virions with antibodies directed against this peptide blocked infection^18^.

For many years, HCMV gB and gMgN were considered to promote initial virion attachment by interacting with cell surface HSPGs. This was mainly based on experiments which studied binding of purified gB and gMgN glycoproteins to heparin columns^23,38^. Yet, infection studies using a gB deletion mutant^39^ or gN-truncated HCMV^40^ had already questioned a major role of gB or gMgN in attachment to host cells. Interestingly, HCMV glycoproteins gH, gL, gO and gB co-expressed in transfected cells have already been shown to promote cell fusion by binding 3-*O*-sulfated heparan sulfate, yet, without revealing the exact heparan sulfate-binding partners^41^. We have shown before that gHgLgO and gB do form complexes in virions^14^ which raised the question whether gHgLgO alone or gHgLgO - gB complexes promote HSPG-binding. Here, we could demonstrate that gHgLgO is the major viral binding partner for cell surface HSPG and that the gO249 mutation abolishes this interaction. Additionally, the gO249 mutation seems to enhance the interaction with gB which very likely additionally blocks gB-dependent binding to HSPGs. Still, gB showed binding to HSPGs independently of gHgLgO albeit to a much lower extent than gHgLgO. This becomes clearly apparent in gO-negative virions.

All experiments presented here show that gHgLgO-dependent attachment to HSPGs is crucial for cell free infection. Yet, it has been shown by us and others that ΔgO mutants show efficient focal spread^9,14^ which is also true for the gO249 mutant^13^. This discrepancy has also been observed *in vivo* in the MCMV infection of mice^19^. Deletion of the MCMV gHgLgO complex abolished infection of mice by preventing infection of first target cells after intravenous application of virus. In contrast, focal spread in for example livers could equally be secured by gHgLgO or the alternative complex gHgLMCK2.

In summary, trimer-dependent tethering of HCMV to HSPGs is a major driver of cell-free infection in cell culture and supposedly also of infection of first target cells *in vivo.* Trimer-dependent tethering is followed by firm docking of trimer or pentamer to their respective entry receptors. Thus, both, tethering and entry receptor binding in concert secure establishment of CMV infections. This suggests that complementation of gB and/or pentamer vaccines with trimer vaccines might be an option to design more efficient HCMV vaccines. For that, future research will have to expand recent studies^42^ on important antibody target sites within the gHgLgO complex.

## Materials and methods

### Cells and viruses

Primary human foreskin fibroblasts (HFF; PromoCell) and HEK293T cells (ATCC: CRL-3216) were maintained in DMEM (Gibco) supplemented with 10% FCS and penicillin/streptomycin. Telomerase-immortalized human microvascular endothelial cells^43^ (TIME cells) were cultured in Endothelial Cell Growth Medium MV 2 (PromoCell). All viruses used are derived from HCMV strain TB40/E cloned as a bacterial artificial chromosome^44^ (TB40-BAC4) and express a firefly luciferase reporter^8^ (TB40-BAC4-luc virus). TB40-BAC4-luc-131stop virus (131stop)^8^ lacks the gHgLpUL(128,130,131A) complex and TB40-BAC4-luc-ΔgO virus^8^ lacks the gHgLgO complex.

### Antibodies, recombinant proteins and reagents

Antibodies specific for HCMV were mouse anti-MCP, human anti-gB SM5-1, mouse anti-gH SA4 (all kindly provided by M. Mach, University Erlangen-Nürnberg, Germany), mouse anti-gH 14-4b (kindly provided by W. Britt, University of Alabama, Birmingham, USA), mouse anti-gO^45^, mouse anti-UL128 4B10 (kindly provided by T. Shenk, University of Princeton, USA), mouse anti-gB 2F12 (Biozol) and rabbit anti-gL LS-C371267 (LSBio). Antibodies specific for cellular proteins or protein tags were rabbit anti-PDGFRα (Cell Signaling Technology), mouse anti-GAPDH GA1R (Thermo Fisher Scientific) and mouse anti-StrepMAB-Classic (IBA Lifesciences GmbH). Secondary antibodies were peroxidase-coupled goat anti-mouse (Sigma), kappa light chain-specific goat anti-mouse (Jackson ImmunoResearch), goat anti-rabbit (Dianova) and goat anti-human (Jackson ImmunoResearch). Soluble recombinant human PDGFRα-Fc fusion protein (R&D Systems) and heparin sodium salt (Sigma) were used for competition experiments.

### Expression and purification of recombinant HCMV gH_Strep_gLgO_WT_

Synthetic genes encoding HCMV gH, gL and gO (Twist Biosciences) were cloned into an insect cell expression vector described previously^46^. To allow for efficient purification, a double Strep-tag was fused to the C-terminus of gH together with an enterokinase (EK) cleavage site facilitating controlled proteolytic cleavage and co-transfected in equimolar ratio into Drosophila S2 cells. To initiate soluble protein expression, the stable cell line was scaled up and induced with 4 μM CdCl_2_ at a cell density of 6×10^6^ cells/mL. 5 days after induction, cells were harvested by centrifugation, and soluble proteins were purified from the supernatant using affinity chromatography on a Strep-Tactin XT 4Flow column (IBA Lifesciences), followed by size-exclusion chromatography using a Superose 6 Increase column (GE Healthcare) equilibrated in PBS. Protein was concentrated to approximately 3.9 mg/ml.

### BAC mutagenesis and virus reconstitution

Charge cluster to alanine (CCTA) mutants gO117 and gO249 were cloned based on a two-step mediated red recombination method^47^. The CCTA mutants were introduced into TB40-BAC4-luc and TB40-BAC4-luc-131stop. Specifically, for the gO117 mutant amino acids **RK**PA**K** at position 117 to 121 were changed to **AA**PA**A** using the primers gO117 forward (5’- TAACCTATCTGTGGTTCGATTTTTATAGTACCCAGCTT**GCTGC**ACCCGCC**GC**ATACGTCT ACTCACAGTACAGGATGACGACGATAAGTAGGG-3’) and gO117 reverse (5’- ATCGTTTTAGCCGTATGATTGTACTGTGAGTAGACGTAT**GC**GGCGGGT**GCAGC**AAGCTG GGTACTATAAAACAACCAATTAACCAATTCTGATTAG-3’) and for the gO249 mutant the amino acids **RK**L**KRK** at position 249 to 254 were changed to **AA**L**AAA** using the primers gO249 forward (5’- CCCCAAGTATATTAACGGCACCAAGTTGAAAAACACTATG**GCTGC**ACTA**GC**A**GC**T**GC**AC AAGCGCCAGTCAAAGAACAAGGATGACGACGATAAGTAGGG-3’) and gO249 reverse (5’- TTTTAGTCTTTTTTTCTAATTGTTCTTTGACTGGCGCTTGT**GC**A**GC**T**GC**TAGT**GCAGC**CA TAGTGTTTTTCAACTTGGCAACCAATTAACCAATTCTGATTAG-3’). Successful mutagenesis was confirmed by restriction fragment analysis and sequencing. To reconstitute BAC DNA to infectious virus, HFF were transfected with 1.5 µg purified DNA using FuGENE HD transfection reagent (Roche) according to the manufacturer’s protocol. Transfected cells were maintained until a strong cytopathic effect (CPE) became visible. To prepare cell-free virus, supernatants were precleared for 15 min at 2600 x g.

### Infections, virus stock preparation, virus titration and growth curves

For infection, medium of 90% confluent HFF or TIME cells was replaced by virus diluted in DMEM containing 5% FCS followed by incubation for 2 h at 37°C. Infection with low titer virus preparations was enhanced by a centrifugation step for 30 min at 860 x g at room temperature (RT) followed by incubation for 1.5 h at 37°C.

For virus stock production, supernatants from infected HFF showing complete CPE were collected and precleared for 15 min at 2600 x g. After preclear, supernatant virus was pelleted by ultracentrifugation for 70 min at 37.700 x g at 4°C, the virus pellets resuspended in DMEM with 5% FCS or VSB buffer (50 mM Tris-HCl, 12 mM KCl, 5 mM EDTA, pH 7.8) and the concentrated stocks stored at −80°C.

Virus titers were determined by a TCID50 assay on HFF.

Multistep growth curves on HFF were performed on 24-well plates. For comparison of different virus mutants, cells were infected such that 24 h post infection, the percentage of initially infected cells was comparable for all viruses to be analyzed. Every second day, the medium was exchanged, the collected supernatants precleared and the virus in supernatants titrated or quantified using qPCR.

### Plasmids and transfection

HCMV glycoproteins expressed from eukaryotic expression vectors are based on the TB40-BAC4 sequence^44^. Full-length ORFs of gH, gL and gB were cloned in the pCR3 vector (Invitrogen). Wildtype gO and gO_249_ were cloned in the pFUSE-mIgG2B-Fc2 vector (InvivoGen). For that, the N-terminal signal peptides (amino acids 1 to 29) of gO_WT_ or gO_249_ were exchanged for the 20 amino acids of the human IL2 signal sequence and the Fc sequence of the vector was deleted. The IL2-gO fusion proteins are thus comprised of amino acids 30 to 464 of HCMV gO. HEK293T cells were transiently transfected on 6-well plates with totally 6 µg DNA mixed with polyethyleneimine (Sigma). 48 h post transfection, cells were lysed for analysis of protein expression by Western blot or for immunoprecipitation experiments.

### Immunoprecipitation

Virions, transfected or infected cells or virus-cell co-incubations were lysed in standard lysis buffer (20 mM Tris-HCl, 150 mM NaCl, 1% Triton X-100, pH 8) containing cOmplete Mini protease inhibitor cocktail (Roche) for 1 h at 4°C. Lysates were cleared for 10 min at 18.000 x g at 4°C and then co-incubated with antibodies, recombinant PDGFRα-Fc or heparin agarose (Sigma) overnight at 4°C. The next day, Protein A Sepharose CL-4B (Avantor) was added to PDGFRα-Fc- or antibody-dependent co-incubations for another 4 h at 4°C. Beads were washed with lysis buffer and then 2x sample buffer (130 mM Tris-HCl, 10% glycerol, 10% 1-thioglycerol, 6% SDS, pH 6.8) was added to dissociate proteins from the beads. Alternatively, the precipitates were digested with sequencing grade modified porcine trypsin (Promega) overnight at 37°C followed by treatment with dithiothreitol (1 mM) for 10 min at 25°C, iodoacetamide (1.35 mM) for 30 min at 25°C and trifluoracetic acid (1.25%). The lyophilized samples were then analyzed by LC-MS/MS.

### Liquid chromatography tandem mass spectrometry (LC-MS/MS) analysis

LC-MS/MS analysis was performed using an Ultimate 3000 nano-liquid chromatography system (Thermo Fisher Scientific) coupled to a Q Exactive HF-X mass spectrometer (Thermo Fisher Scientific). Peptides were diluted in 0.1% formic acid and transferred to an Acclaim PepMap 100 trap column (nanoViper C18, 2 cm length, 100 μM ID, Thermo Fisher Scientific). An analytical EasySpray column (PepMap RSLC C18, 50 cm length, 75 μm ID, Thermo Fisher Scientific) was used for separation. Liquid chromatography was performed at a flow rate of 250 nl/min using 0.1% formic acid as solvent A and 0.1% formic acid in acetonitrile as solvent B. Peptides were eluted using an 80 min gradient from 5% to 20% solvent B, followed by a 10 min gradient from 20% to 40% B. Eluting peptides were analyzed using a data-independent acquisition method with 50×12 m/z wide precursor isolation windows in the range of 400-1000 m/z in a staggered window pattern. MS RAW data were processed using DIA-NN (v1.8)^48^ with spectral libraries generated in DIA-NN by deep learning-based spectra and retention time prediction. As input sequence databases, the human subset of the UniProt database and a database consisting of the following HCMV proteins based on TB40-BAC4 were used: gH, gL, gO_WT_, gO_249_, UL128, UL130, UL131A and gB. In addition, the sequence of *Mus musculus* Ighg2b (UniProt: P01867) was implemented to allow quantification of the antibody as loading control of the immunoprecipitates. Volcano plot analysis was performed using Perseus (1.5.3.2)^49^ with an unpaired t-test and default values (FDR = 0.05, S0=0.1) as significance threshold. The mass spectrometry proteomics data have been deposited to the ProteomeXchange Consortium via the PRIDE^50^ partner repository with the dataset identifier PXD058164.

### Western blot analysis

To analyze proteins in virus particles, cells or immunoprecipitates by WB, samples were lysed in 2x sample buffer. Proteins were subjected to SDS-polyacrylamide gel electrophoresis followed by WB analysis using nitrocellulose membranes for transfer of the proteins (0.45 µm, Thermo Fisher Scientific). Membranes were blocked with 5% skim milk and then incubated with primary and secondary antibodies. Antibody binding was detected using Super Signal West Pico chemiluminescence substrate (Thermo Fisher Scientific).

### Virion DNA isolation and qPCR

For viral DNA isolation from concentrated virus stocks or cell culture supernatants, DNA not protected in viral capsids was digested with 4 U DNAse I (NEB) for 1.5 h at 37°C. Then, 5 mM EDTA was added and the enzyme was heat-inactivated for 10 min at 75°C.

Viral DNA in particles was isolated using the DNeasy Blood & Tissue Kit (QIAGEN) according to the manufacturer’s protocol. Briefly, the DNAse I-digested samples or virions attached to cells were incubated with Proteinase K in lysis buffer for 10 min at 56°C followed by the addition of ethanol. Then, the samples were transferred onto the provided columns, washed and the extracted DNA was eluted in DNAse and RNAse free ddH_2_O.

To quantify the amount of virion DNA, qPCR was performed. A reaction mixture of 20 µl containing 1x SYBR Green PCR Master Mix (Applied Biosystems), 2 µl isolated virion DNA and 500 nM of UL83 forward (5’-TGGTCACCTATCACCTGCAT-3’) and reverse (5’- GAAAGAGCCCGACGTCTACT-3’) primer was prepared. DNA was detected using the QuantStudio 5 Real-Time-PCR-System (Applied Biosystems). To calculate the number of viral genomes (copies/ml), a TB40-BAC4 bacmid standard was used.

### Heparinase digestion

To remove cell surface HSPGs, HFF or TIME cells were incubated for 2 h at 30°C with heparinase I, II and III (1 U/ml, NEB) in Opti-PRO medium (Gibco) supplemented with 25 mM HEPES (Gibco). Excess heparinases were removed by washing the cell monolayers with DPBS.

### Luciferase assay

Luciferase assays were performed as described previously^8^. Briefly, HFF or TIME cells were seeded on 96-well plates and infected the next day. If inhibition assays were performed, PDGFRα-Fc or heparin were serially diluted and mixed with the respective viruses. For PDGFRα-Fc inhibition, the virus – PDGFRα mixtures were pre-incubated for 1 h at 4°C before virus was added. For infection, cells and viruses were co-incubated for 1.5 h at 37°C. Then, the supernatants were exchanged for medium containing phosphonoacetic acid (300 µg/ml, Sigma). 48 h post-infection, the luciferase activity in infected cells was determined using the Firefly Luciferase Assay Kit 2.0 (Biotium) according to the manufacturer’s protocol.

### Binding assay

HFF or TIME cells were seeded in 24-well plates. The next day, the cells were pre-cooled for 10 min on ice before virus particles were added in a volume of 250 µl medium. To inhibit binding to HSPGs, heparin (100 µg/ml) was added to the medium or the cells were pre-incubated with heparinases before addition of virus. Binding was allowed for 1 h on ice followed by three washing steps at 4°C to remove unbound virus. Finally, DNA was extracted from the cell monolayers using the DNeasy Blood & Tissue Kit and qPCR was performed to determine the number of virion particles bound to the cells.

### ELISA for gH_strep_gLgO quantification

HFF or TIME cells were seeded on 96 well plates. The next day, cells were incubated with recombinant HCMV gH_Strep_gLgO (100 µg/ml) for 1 h at 37°C. After removal of unbound protein, the cells were fixed with 1% PFA for 10 min at RT. To detect gH_Strep_gLgO_WT_ complexes bound to cell surfaces, cells were incubated with mouse anti-Strep antibody for 1 h at RT followed by incubation with peroxidase-coupled goat anti-mouse antibody (1:2000) for 1 h at RT. Then, TMB substrate (BioLegend) was added onto the cells. The reaction was stopped with 10% phosphoric acid and the absorbance determined at 450 nm.

### Data visualization, analysis and statistics

Data were analyzed and statistical significance was determined with GraphPad Prism 10.1.2. Structures were visualized using PyMOL 2.5.5. Schematic presentations were created with BioRender and Western blots were quantified with ImageJ 1.52a.

## Acknowledgements

We thank Heiko Adler for critically reading the manuscript. BA was supported by a grant from the German Center for Infection Research (TTU 07.833).

## Author contributions

B.A. acquired funding. B.A. and L.T. conceptualized the project, performed the data analysis and wrote the manuscript. L.T. curated the data. L.T., R.G., L.K. and B.A. performed the experiments. T.F. and M.K. performed the LC-MS/MS analysis. I.G. constructed the pFUSE-gO expression plasmid. D.A.S. and T.K. cloned expression vectors for insect cells and purified recombinant gHgLgO.

