## Supplementary Material for "A dual role in virion attachment and entry makes the human cytomegalovirus gHgLgO trimer the central player in virion infectivity"

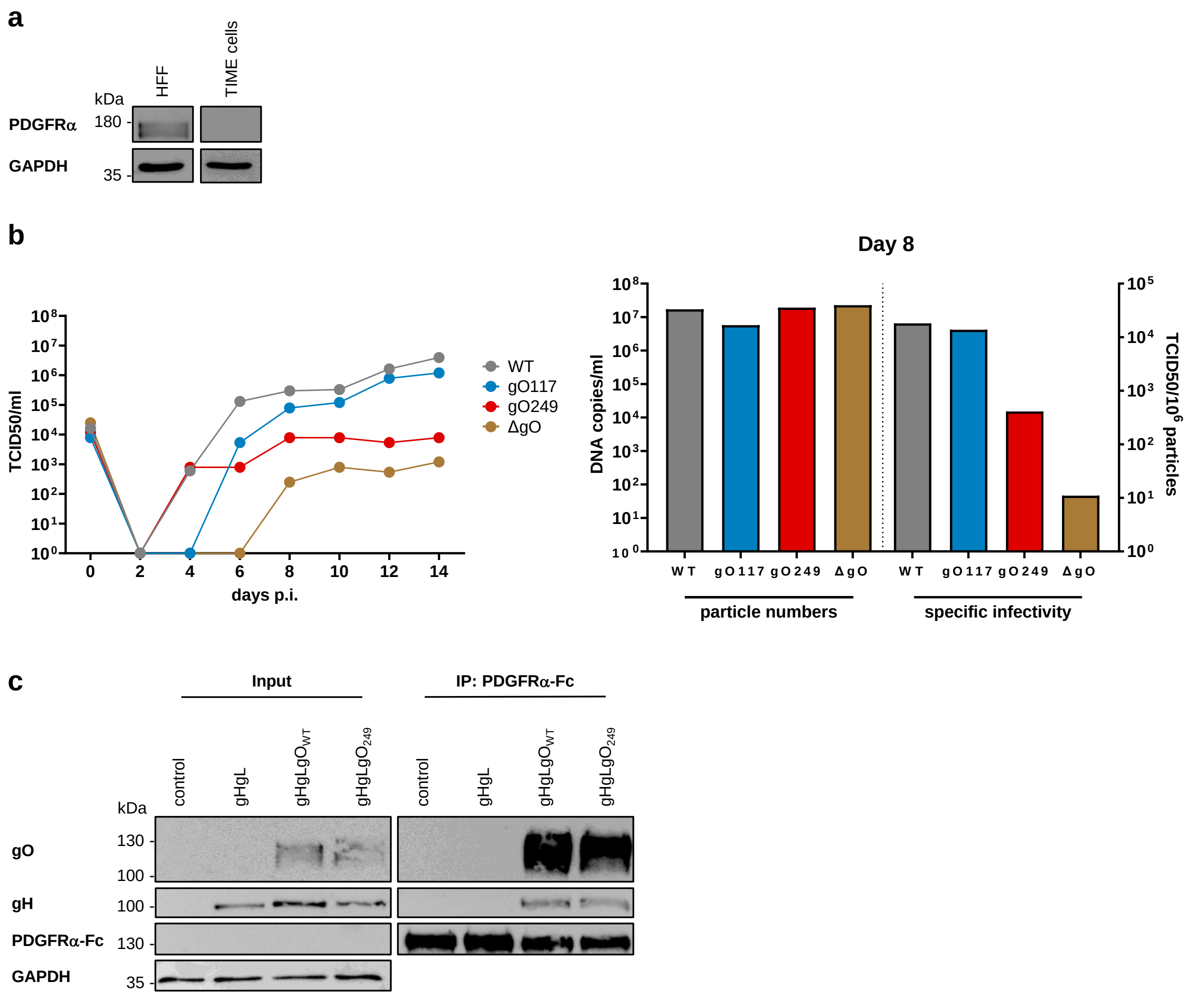


**Supplementary Figure 1. Complementary characterization of cells and gO mutants**

(**a**) Endogenous PDGFRα expression levels in HFF and TIME cells were determined by WB analysis of total cell lysates. GAPDH served as a loading control. (**b**) Multistep growth curves of different gO mutants on HFF. Infectious supernatant virus was determined by a TCID50 assay. Right panel: Particle numbers quantified by qPCR and specific infectivity of virus particles determined for day eight growth curve supernatants. (**c**) WB analysis of gH and gO precipitated from lysates of HEK293T cells expressing GFP (control), gHgL, gHgLgO$_{WT}$ or gHgLgO$_{249}$ using recombinant PDGFRα-Fc. GAPDH levels served as loading control. (**a**, **b** and **c**) Shown are representative experiments.

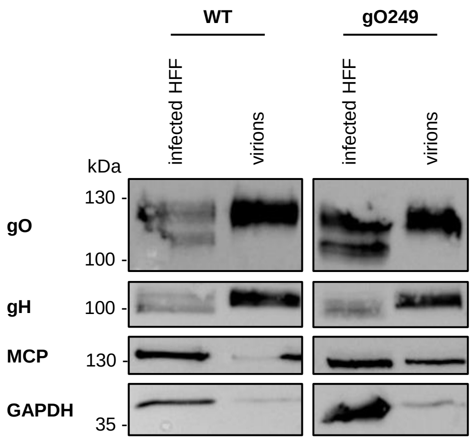


**Supplementary Figure 2. Analysis of gH and gO expressed in infected HFF and virions**
WB analysis of gH and gO in lysates of HFF infected with WT or gO249 virus and in lysates of the respective virions. MCP and GAPDH expression levels served as loading controls. One representative experiment is shown.

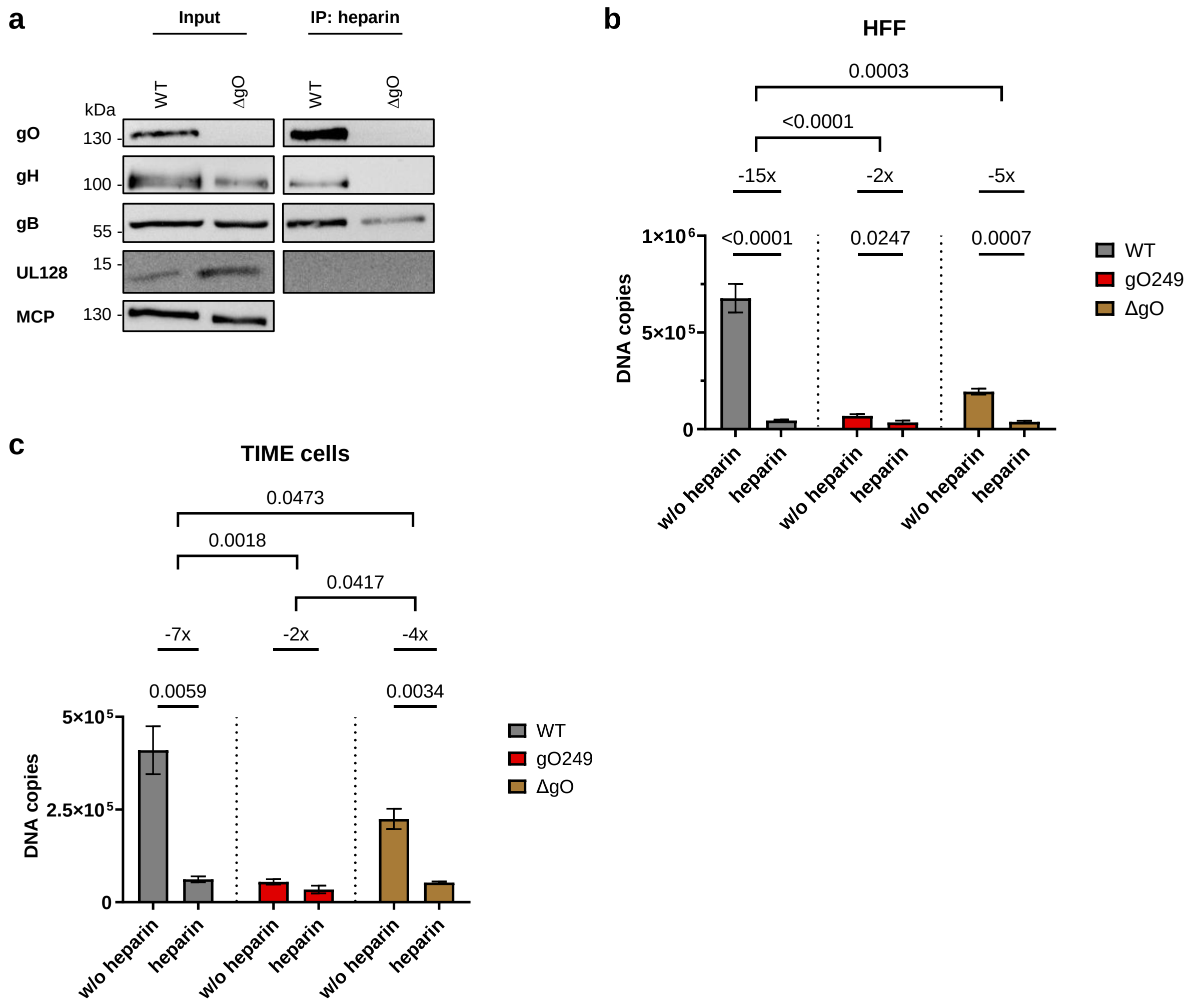


**Supplementary Figure 3. A role of gB in HSPG-dependent tethering**

(**a**) WB analysis of gH, gO, gB and UL128 precipitated from lysates of WT or ΔgO virions using heparin agarose. One representative experiment of four is shown. (**b**) HFF or (**c**) TIME cells were co-incubated with $5x10^7$ WT, gO249 or ΔgO virus particles in the presence (100µg/ml) or absence of heparin and virus particles bound to cells were quantified by qPCR. Shown are means +/- SEM of independent experiments ($n$ = 3-6). Statistical significance was determined for pairwise comparisons of heparin-treated or -untreated infections (Student's unpaired t-test). Additionally, fold changes were calculated and analyzed (one-way ANOVA). $P$ values of statistically significant differences are depicted. The data showing virus attachment of WT and gO249 virions both in the presence and absence of heparin are identical to the data of Figure 3e and f.

| | Protein | Fold change | P value |
|---|---|---|---|
| HCMV glycoproteins | Envelope glycoprotein L | 0.012 | 0.00009 |
| | Envelope glycoprotein O | 0.018 | 0.00022 |
| | Envelope glycoprotein H | 0.025 | 0.00018 |
| | Envelope glycoprotein B | 0.035 | 0.00042 |
| Human proteins | Trifunctional enzyme subunit beta, mitochondrial | 0.029 | 0.00001 |
| | Syndecan-1 | 0.033 | 0.00018 |
| | Syndecan-4 | 0.038 | 0.00024 |
| | Probable glutathione peroxidase 8 | 0.057 | 0.00083 |
| | Vesicle-associated membrane protein 3 | 0.060 | 0.00024 |
| | Exocyst complex component 3 | 0.063 | 0.00010 |
| | Fibroblast growth factor 2 | 0.066 | 0.00883 |
| | Trifunctional enzyme subunit alpha, mitochondrial | 0.071 | 0.00231 |
| | Lipase maturation factor 2 | 0.075 | 0.00023 |
| | ATP-binding cassette sub-family D member 3 | 0.075 | 0.00012 |
| | Protein RER1 | 0.077 | 0.00078 |
| | Neuropilin-1 | 0.079 | 0.00002 |
| | Protein transport protein Sec61 subunit alpha isoform 1 | 0.093 | 0.00228 |
| | Exocyst complex component 2 | 0.101 | 0.00800 |
| | Exocyst complex component 4 | 0.102 | 0.00085 |
| | Exocyst complex component 1 | 0.102 | 0.00131 |
| | Transforming growth factor beta receptor type 3 | 0.117 | 0.00434 |
| | Oxysterol-binding protein-related protein 8 | 0.119 | 0.00005 |
| | Transmembrane protein 109 | 0.138 | 0.01113 |
| | DNA-dependent protein kinase catalytic subunit | 0.143 | 0.02362 |
| | Syndecan-2 | 0.147 | 0.01848 |
| | Wolframin | 0.148 | 0.00153 |
| | Histone-lysine N-methyltransferase 2B | 0.152 | 0.00306 |
| | Vitamin K epoxide reductase complex subunit 1 | 0.153 | 0.00688 |
| | Solute carrier family 25 member 3 | 0.174 | 0.00164 |
| | Sequestosome-1 | 0.186 | 0.00328 |
| | Sorting nexin-18 | 0.191 | 0.00557 |
| | Platelet-derived growth factor receptor alpha | 0.196 | 0.00407 |
| | Very-long-chain enoyl-CoA reductase | 0.197 | 0.00198 |
| | E3 ubiquitin-protein ligase Itchy homolog | 0.201 | 0.01019 |
| | Glypican-6 | 0.202 | 0.00113 |
| | CD44 antigen | 0.207 | 0.00159 |
| | Surfeit locus protein 4 | 0.213 | 0.01988 |
| | Tricarboxylate transport protein, mitochondrial | 0.225 | 0.01664 |
| | Protein S100-A6 | 0.243 | 0.00094 |
| | Matrix metalloproteinase-14 | 0.257 | 0.00192 |
| | UDP-glucose 6-dehydrogenase | 0.271 | 0.01957 |
| | Caveolin-1 | 0.301 | 0.00669 |
| | Voltage-dependent anion-selective channel protein 2 | 0.333 | 0.00304 |
| | Small ribosomal subunit protein uS7 | 0.380 | 0.00900 |
| | ADP/ATP translocase 2 | 0.425 | 0.00366 |
| | Ras-related protein R-Ras | 0.436 | 0.00400 |
| | Transmembrane protein 43 | 0.455 | 0.01261 |
| | ADP-ribosylation factor 4 | 0.457 | 0.00744 |
| | Eukaryotic initiation factor 4A-I | 0.522 | 0.00093 |
| | EH domain-containing protein 2 | 0.565 | 0.00100 |
| | Aspartate-tRNA ligase, cytoplasmic | 0.675 | 0.00017 |

**Supplementary Table 1. LC-MS/MS hits of interaction partners of 131stop-gO249 versus 131stop virions**

LC-MS/MS data from anti-gH (14-4b) immunoprecipitates of lysates of HFF co-incubated with 131stop or 131stop-gO249 virions. Data correspond to the Volcano plot shown in Figure 2c and are depicted as fold change 131stop-gO249 virus versus 131stop virus. Additionally, the respective *P* values are shown. Selected proteins and proteoglycans are highlighted (HCMV glycoproteins (red), PDGFR$\alpha$ and TGF$\beta$RIII (green), HSPGs (blue)).
